# A dual role for peripheral GDNF signaling in nociception and cardiovascular reflexes

**DOI:** 10.1101/627521

**Authors:** Luis F. Queme, Alex A. Weyler, Elysia R. Cohen, Renita C Hudgins, Michael P. Jankowski

**Affiliations:** Department of Anesthesia, Division of Pain Management, Cincinnati Children’s Medical Center, University of Cincinnati, College of Medicine, Cincinnati OH 45229; Department of Pediatrics, University of Cincinnati, College of Medicine, Cincinnati OH 45229

## Abstract

Group III/IV muscle afferents transduce nociceptive signals and modulate exercise pressor reflexes (EPR). However, the mechanisms governing afferent responsiveness to dually modulate these processes are not well characterized. We and others have shown that ischemic injury can induce both nociception-related behaviors and exacerbated EPRs in the same mice. This correlated with primary muscle afferent sensitization and increased expression of glial cell line-derived neurotrophic factor (GDNF) in injured muscle and increased expression of GDNF family receptor α1 (GFRα1) in DRGs. Here we report that increased GDNF/GFRα1 signaling to sensory neurons from ischemia/reperfusion affected muscle modulated nociceptive-like behaviors, increased EPRs, and group III/IV muscle afferent sensitization. This appeared to have taken effect through increased CREB/CREB-binding protein mediated expression of the purinergic receptor P2X5 in the DRGs. Muscle GDNF signaling to neurons may play an important dual role in nociception and sympathetic reflexes and could provide a novel therapeutic target for treating complications from ischemic injuries.

## Introduction

Group III and IV primary muscle afferents are subpopulations of thinly myelinated (III) and unmyelinated (IV) fibers with a variety of sensory capabilities that include mechanical-, thermal-, chemo-sensation ^1, 2, 3^. Group III/IV muscle afferents have two distinct functions. First, these neurons can serve as sensory transducers of noxious and non-noxious peripheral stimuli from the muscles ^2, 4^. Second, they also function as the sensory arm for the exercise pressor reflex (EPR), the cardiovascular response to muscle contraction that includes increased blood pressure and heart rate ^5, 6^. Each of these biological processes are influenced by the different sensory modalities of group III/IV afferents ^7, 8, 9, 10, 11, 12^.

A common method to study the involvement of peripheral sensory neurons in both nociception and sympathetic reflexes is to use a skeletal muscle ischemic injury model ^13, 14, 15, 16, 17^. After ischemic injuries, group III and IV muscle afferents display peripheral sensitization, including increased responsiveness to mechanical and chemical stimuli. Ischemia also increases the response to muscle contractions of ∼50% of group IV and ∼12% of group III muscle afferents ^14^. This correlates with observed changes in nociceptive behaviors, EPRs, and gene expression in the affected muscles and dorsal root ganglia (DRGs) ^7, 13, 18^.

Chemo-reception and mechanical responsiveness of primary muscle afferents have been attributed to expression of a combination of both acid sensing ion channels (ASICs) and purinergic, P2X receptors ^7, 19, 20, 21, 22, 23^. Each of these ion channels have also been associated with altered nociception and EPR modulation after ischemic injury ^8, 16, 24, 25^. P2X and ASIC channel activity has been linked to sensing of fatigue and ischemic pain ^7, 19^ by disrupting the metabolite responses of DRG neurons ^7, 9, 19^. P2X5 has also been shown to modulate the pH sensitivity of ASIC3 ^19^, which may further shape afferent and behavioral responsiveness. The role of these channels in the normal mechanical responses of nociceptors is not as clear, but both ASIC3 and P2X receptors have been linked to mechanical hypersensitivity in different injury models ^7, 16, 20, 23, 26^.

We previously reported an important role for the cytokine interleukin 1-beta (IL-1β) modulating the expression of ASIC3 in mechanically and chemically sensitive afferents in the DRGs to regulate peripheral sensitization after ischemia with reperfusion (I/R) injury ^16^. This suggested that factors expressed in the injured muscles may have a substantial influence on afferent function and subsequent behaviors after I/R. In addition to studies on cytokine production, we and others also recently reported that growth factors may be additional signaling molecules involved in modulating muscle injury-related hypersensitivity ^27, 28, 29, 30^. One of the striking findings of our work was that of all of the numerous growth factors (GF) tested, only Glial Cell-Derived Neurotrophic Factor (GDNF) was found to be upregulated in the muscle one day after I/R injury. Its receptor, GDNF family receptor α1 (GFRα1), was also specifically induced in the I/R affected DRGs ^15^. Previous studies have suggested that GDNF can sensitize peripheral nociceptors ^31^, and it has been reported that local injection of GDNF is capable of producing muscle mechanical hypersensitivity ^32^. Furthermore, intramuscular injection of anti-GDNF targeting antibodies have been shown to prevent the development of mechanical hypersensitivity in a rat model of delayed onset muscle soreness ^33^. GDNF has been associated with increased mechanical responsiveness in muscle afferents ^34^, and GDNF can regulate sensory neurons expressing both ASIC3 and P2X receptors ^31, 35^.

Clinically, deep tissue I/R injuries are common characteristics of multiple conditions, such as peripheral arterial disease (PVD) and sickle cell anemia. A common hallmark of these diseases is chronic musculoskeletal pain that is often observed alongside an exacerbation of the patient’s EPR ^36, 37^. In clinical settings, the changes associated with muscle ischemia can cause pain and lead to exercise intolerance or further complicate underlying cardiovascular conditions ^13, 37, 38, 39^. While there is a plethora of knowledge regarding EPRs and the peripheral mechanisms of nociception from the skin ^31, 40, 41^, there is a significant disparity in our understanding of afferent mechanisms of myalgia, especially in the context of ischemia. Further, whether peripheral mechanisms of muscle nociception share common pathways with mechanisms of EPR modulation is relatively unexplored ^13, 42^. Here we hypothesized that the GDNF/GFRα1 pathway sensitizes primary muscle afferents after ischemic insults and regulates the observed increase in the cardiovascular response to exercise as well as pain-related behaviors.

## Results

### I/R upregulates GDNF in the affected muscle and various gene expression changes in the DRGs that correlate with behavioral hypersensitivity

In our previous reports, we did not observe a significant increase in the expression of many growth factors in the I/R affected muscle tissue such as NGF, BDNF and Artemin. However, GDNF was significantly upregulated ^15^. To therefore characterize the role of GDNF signaling after I/R, we first quantified the amount of GDNF protein in muscle tissue by Western Blot. This analysis revealed an increased amount of GDNF in the I/R injured muscles compared to naïve mice 24h after injury (Fig. 1A). Immunohistochemical labeling experiments revealed expression of GDNF around and within myofibers (Fig 1B).

**Figure 1.**
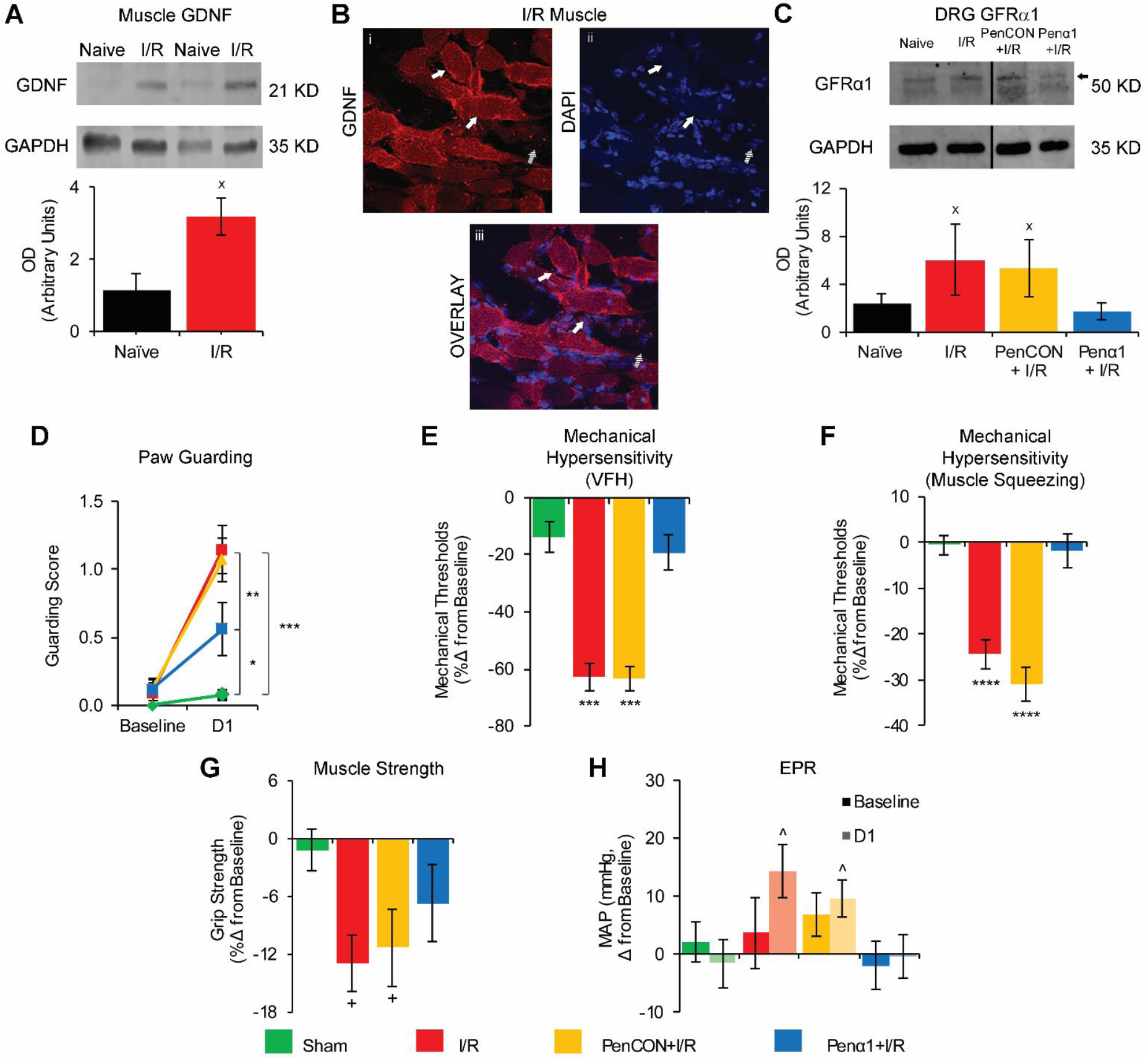
Increased expression of GDNF in muscle and upregulation of GFRα1 in the DRG modulate pain related behaviors and increased mean arterial pressure (MAP) after I/R. **A**. GDNF is increased in muscle after I/R compared to naïve animals (n=4 per group) as assessed by western blotting (WB). **B**. Representative image of GDNF distribution within myofibers in an injured forepaw muscle. DAPI was used to mark nuclei. Solid arrows: GDNF positive myofibers. Dashed arrows: GDNF negative myofiber. **C**. WB of GFRα1 in DRGs showed increased expression in the C7-T1 DRGs after I/R and in PenCON+I/R groups. Penα1 effectively prevented this upregulation (n=3 per group). **D**. Paw guarding is increased 1d after I/R and PenCON+I/R and this is partially prevented in Penα1+I/R animals. Mechanical withdrawal thresholds to von Frey hair (VFH) stimulation (**E**) and muscle squeezing (**F**) are decreased after I/R and PenCON+I/R. GFRα1 knockdown blocked these effects. **G**. Grip strength is decreased 1d after I/R and PenCON+I/R. This was partially reversed in Penα1+I/R animals. **H**. Only injured mice (I/R and PenCON+I/R) displayed a significant increase in MAP after forced exercise. The increase in MAP was not observed in Penα1+I/R mice. (D-H: n=12 per group. ^x^p<0.5 vs Naïve, *p<0.5 vs Sham, **p<0.01 vs Penα1+I/R, ***p<0.001 vs Sham and Penα1+I/R, ****p<0.0001 vs Sham and Penα1+I/R, ^+^p<0.01 vs Sham, ^^^p<0.05 vs pre-exercise MAP; 1-way ANOVA with HSD post hoc test (A, C) or 2-way RM ANOVA with HSD post hoc test (D-H)).

While GDNF showed increased expression in the I/R-injured muscle, its receptor, GFRα1, was upregulated in the I/R-affected DRGs (Fig 1C). In order to test the specific effects of GFRα1 upregulation in DRG neurons, we used our previously described nerve-specific siRNA-mediated knockdown strategy to block dynamic injury-related gene expression ^13, 16, 41, 43, 44^. We verified the effectiveness of the siRNA knockdown strategy via realtime PCR and Western Blot. To control for the nerve injection, a separate group of I/R-injured mice was injected with non-targeting siRNAs (PenCON+I/R). The PenCON+I/R mice showed the same increased mRNA expression of GFRα1 (438%±32%; p<0.05 vs. naive) as the I/R mice without siRNA injections (236%±17%; p<0.05 vs naive), while the Penα1+I/R mice (0%±19%; p>0.05 vs naïve; 1-way ANOVA with HSD post hoc) showed expression levels similar to the naïve animals. Comparable results were also obtained at the protein level (Fig 1C).

We then performed a variety of behavioral tests in I/R injured animals with nerve-targeted GFRα1 knockdown. In these experiments, sham surgery control was used for comparisons. I/R-injured and PenCON injected animals with I/R showed increased paw guarding scores compared to sham injured animals at 1d. I/R-injured mice with GFRα1 knockdown (Penα1+I/R) however show significantly reduced paw guarding (Fig 1D). Withdrawal thresholds to von Frey filament stimulation of the plantar surface of the fore paw were also significantly decreased in the I/R and PenCON+I/R groups compared to sham injured animals. This mechanical hypersensitivity was completely prevented by the Penα1 injection (Fig 1E). Similar results regarding mechanical hypersensitivity were also obtained in groups that underwent hind paw muscle squeezing (Fig 1F). Grip strength was decreased after I/R alone and in the PenCON+I/R group. This decrease in grip strength was partially prevented by GFRα1 targeting siRNAs (Fig 1G). Finally, we tested the cardiovascular response to exercise before and following injury to measure blood pressure and heart rate after a low intensity exercise. This running protocol has been previously determined to be sufficient to induce an EPR response but not strong enough that it can induce pain-related hypersensitivity ^13^. I/R-injured and PenCON+I/R mice showed a significant increase in the mean arterial pressure (MAP) one day after injury when compared with their pre-exercise baseline. This significant increase in MAP was completely absent in sham injured or Penα1+I/R mice (Fig 1H). We did not detect any change in the heart rate after exercise either before or 1 day after I/R injury (not shown). Results collectively indicate that afferent GFRα1 upregulation in the DRG is important in dually modulating nociception and EPRs after I/R.

### GFRα1 upregulation modulates primary afferent sensitization after I/R

To test the hypothesis that GFRα1 upregulation played a role in afferent sensitization after I/R, we performed electrophysiological recordings in an *ex-vivo* muscle, nerve, DRG, spinal cord preparation. As previously reported ^15, 16^, we found that primary group III and IV muscle afferents from I/R and PenCON+I/R mice have decreased mechanical thresholds compared to naïve animals. GFRα1 knockdown prevented observed mechanical sensitization in sensory neurons (Fig 2A, 2B). As shown in our previous work ^15, 16^, naïve afferents in the current report were found to respond almost exclusively to either 15 mM lactate, 1 µM ATP, pH 7.0 (“low” metabolites) or 50 mM lactate, 5 µM ATP, pH 6.6 (“high” metabolites), but not both (Fig 2C). I/R and PenCON+I/R groups showed an increase in the total number of afferents that responded to stimulation of the muscles with both “low” and “high” metabolite mixtures. This phenotypic change was prevented by selective GFRα1 knockdown. Penα1+I/R mice not only showed responses exclusively to one or the other concertation of metabolites, but surprisingly the percentage of afferents recorded showing any response to metabolites was lower than that observed in naïve animals (Fig 2C). This suggests that GFRα1 upregulation not only plays a role in afferent mechano-sensation after I/R, but is also important for chemo-sensory functions of group III/IV afferents.

**Figure 2.**
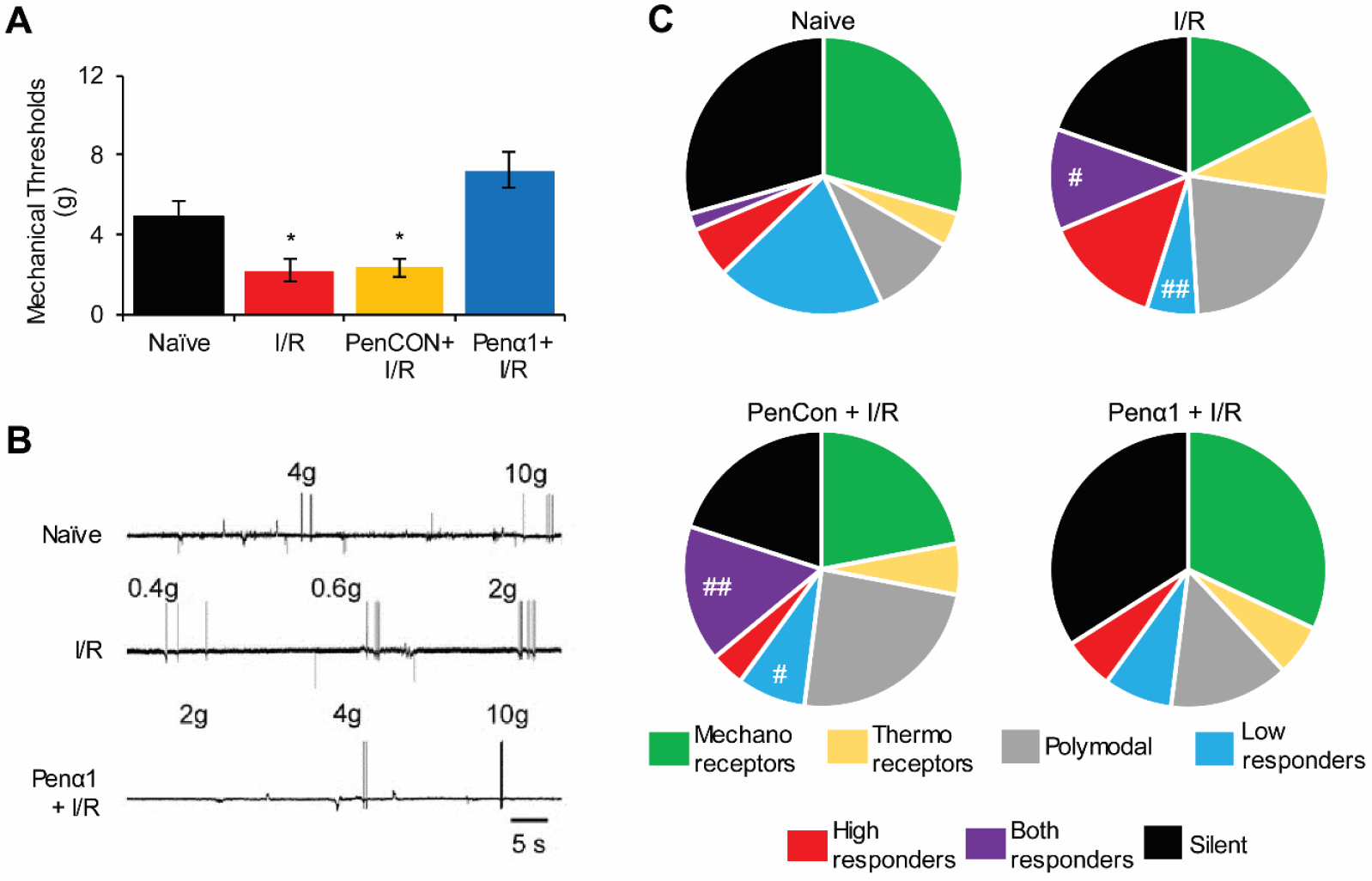
I/R induces mechanical hypersensitivity in group III and IV muscle afferents and a change in the prevalence of chemosensitive fibers. **A**. Primary group III and IV muscle afferents showed decreased mechanical thresholds 1d after I/R alone or PenCON+I/R compared to naïve. Penα1 injection effectively prevented this I/R induced mechanical sensitization (Naïve n=26, I/R n=21, PenCON+I/R n=26, Penα1+I/R n=20). **B**. Representative traces of afferent mechanical responses from select groups. Grams used to elicit responses are indicated above the individual responses. **C.** In I/R and PenCON+I/R groups, there was a significant decrease in the total number of “Low Responders” and a significant increase in the total number of “Both Responders.” These changes were not observed in Penα1+I/R animals (Naïve n=51, I/R n=51, PenCON+I/R n=50, Penα1+I/R n=50; *p<0.05 vs Naïve and Penα1+I/R, ^#^p<0.05 vs Naïve, ^##^p<0.01 vs Naïve; 1-way ANOVA with HSD post hoc (B) and χ2 (C)).

### GFRα1 regulates select changes in DRG gene expression after I/R

The increased expression of GFRα1 in the affected DRG was accompanied by a significant upregulation of various genes encoding receptors involved in sensory transduction. Similar to previous reports ^15, 16^, we found that the acid sensing ion channels 1 (ASIC1) and ASIC3, and purinergic receptors P2X3, P2X4 and P2X5 were significantly upregulated 1d after I/R. Other receptors from the GFR family, including GFRα2 and GFRα3 were not upregulated after I/R. The tyrosine receptor kinase (trk) family of receptors trkA, trkB and trkC were also not upregulated in the DRGs after I/R (Table 1).

**Table 1:** Select DRG gene expression 1d after I/R. Values indicate percent change versus naïve. mean ± SEM. Yellow = p<0.05 vs. naïve, 1-way ANOVA.

| Gene: DRGs | I/R |
| --- | --- |
| ASIC1 | 364 ± 15% |
| ASIC3 | 110 ± 10% |
| P2X3 | 114 ± 10% |
| P2X4 | 194 ± 10% |
| P2X5 | 70 ± 10% |
| GFRα1 | 236 ± 16% |
| TrkA | -23 ± 1% |
| TrkB | -22 ± 1% |
| TrkC | -19 ± 2% |
| GFRα2 | -5 ± 1% |

Percent Change in mRNA

We therefore assessed the effects of GFRα1 knockdown on upregulated receptor expression in the DRGs after I/R. We did not find any significant difference in the expression levels of ASIC1, ASIC3, P2X3, P2X4, P2X5 or GFRα1 between I/R and PenCON+I/R mice (not shown) and thus grouped the data for simplicity of presentation (I/R Control). As shown in Table 2, Penα1+I/R animals showed a significant decrease in the expression level of ASIC3, but not ASIC1 compared to I/R injured animals. However, knockdown did not completely revert levels of ASIC3 to that observed in uninjured mice. Interestingly, the only purinergic channel whose increased expression was significantly blocked by selective GFRα1 knockdown after I/R was P2X5. No I/R induced changes in P2X3 or P2X4 were observed in mice with Penα1 injection plus I/R.

**Table 2:** Effects of GFRα1 knockdown on I/R-related gene expression in DRGs. Values indicate percent change versus naïve. mean ± SEM. n=6 per group, combined I/R Control, n=12. Orange = p<0.001 vs Naïve, Yellow = p<0.01 vs Naïve, Green = p<0.01 vs Naïve and p<0.001 vs I/R Control; 1-way ANOVA with HSD post hoc test.

| Gene:<br>DRGs | I/R<br>Control | Penα1+I/R |
| --- | --- | --- |
| ASIC1 | 511 ± 11% | 520 ± 17% |
| ASIC3 | 112 ± 9% | 48 ± 9% |
| P2X3 | 100 ± 14% | 108 ± 4% |
| P2X4 | 386 ± 17% | 232 ± 10% |
| P2X5 | 69 ± 19% | 18 ± 10% |
| GFRα1 | 438 ± 31% | 0 ± 19% |

These latter results were then corroborated by total cell counts in the DRGs where I/R and PenCON+I/R animals showed a significant increase in the total number of individual cells positive for either GFRα1 or P2X5, and the total number of neurons co-expressing GFRα1 and P2X5 (Fig. 3). Both of these increases in total number of immunopositive cells were prevented by selective knockdown of GFRα1, suggesting a direct relationship between GFRα1 and P2X5 expression after injury.

**Figure 3.**
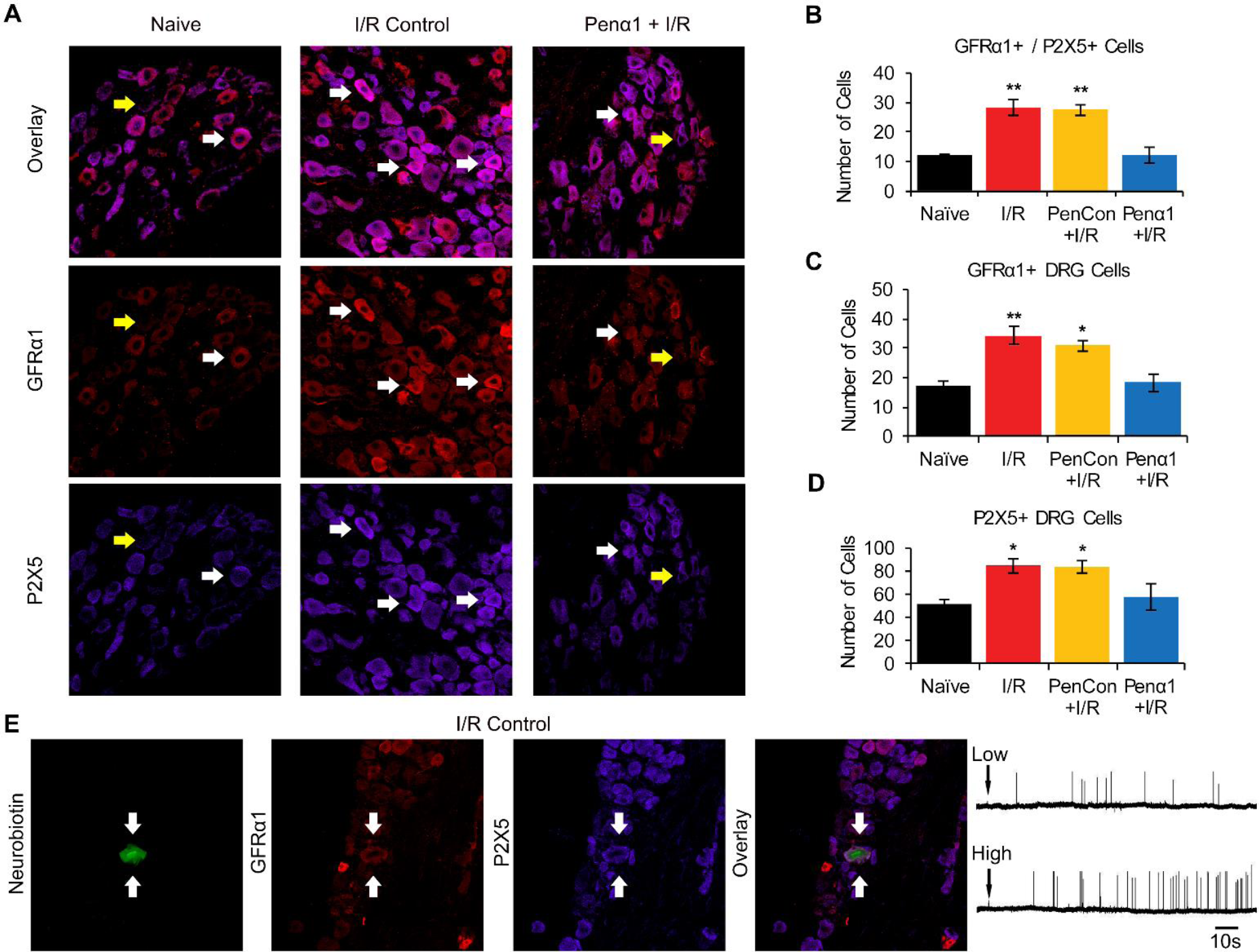
I/R causes and increase in the total number of cells that are positive for GFRα1 and P2X5. **A**. Representative images of DRG sections immunostained for GFRα1 and P2X5. Yellow arrows mark GFRα1 negative cells and white arrows mark positive cells. **B-D**. After I/R there was a significant increase in the total number of cells immunopositive for both GFRα1 and P2X5 **(B)** as well as total cells positive for either GFRα1 **(C)** or P2X5 **(D)**. The increase in immunopositive cells was prevented in Penα1+I/R. (n=3 per group, *p<0.05 vs Naïve and Penα1+I/R, **p<0.01 vs Naïve and Penα1+I/R; 1-way ANOVA with HSD post hoc test). **E.** Representative functionally identified muscle afferent obtained during *ex vivo* recording intracellularly filled with neurobiotin (green) was found to be immunopositive for GFRα1 (red) and P2X5 (blue) and was responsive to both “low” and “high” metabolite mixtures.

To gain better insight on whether the phenotypic alterations in chemosensitive muscle afferents induced by I/R corresponded with the expression of GFRα1 and P2X5, after electrophysiological characterization of identified muscle afferents we filled chemosensitive neurons with neurobiotin and performed immunohistochemistry in the DRG containing the labeled neuron. As shown in Figure 3E and Table 3, after I/R, ∼90% (9/10 GFRα1+, 9/10 P2X5+) the neurons that became responsive to both metabolite mixtures expressed either P2X5 or GFRα1 and 80% of these expressed both receptors (8/10 GFRα1+/P2X5+). In the “low” responder subpopulation, only 25-30% of cells were positive for both receptors (1/4 in Naïve, 1/3 in I/R Control and 0/1 in Penα1+ I/R, Table 3). This suggests that while the co-expression of GFRα1 and P2X5 is not a requirement for the normal chemosensitive function of metaboreceptors, there may be a strong link between the co-expression of both GFRα1 and P2X5, and the phenotypic switch observed in chemosensitive primary muscle afferents after I/R.

**Table 3:** Expression of GFRα1 and P2X5 in chemosensitive group III/IV muscle afferents.

| Receptor Type | Condition | GFRα1+ | P2X5+ | GFRα1+/<br>P2X5+ |
| --- | --- | --- | --- | --- |
| Low Responders | Naïve | 2/4 | 1/4 | 1/4 |
|  | I/R Control | 1/3 | 3/3 | 1/3 |
|  | I/R | 0/1 | 1/1 | 0/1 |
|  | PenCON+I/R | 1/2 | 2/2 | 1/2 |
| | Penα1+I/R | 0/1 | 0/1 | 0/1 |
| High Responders | Naïve | 1/1 | 1/1 | 1/1 |
|  | I/R Control | 3/4 | 3/4 | 3/4 |
|  | I/R | 2/2 | 2/2 | 2/2 |
|  | PenCON+I/R | 1/2 | 1/2 | 1/2 |
| | Penα1+I/R | 0/0 | 0/0 | 0/0 |
| Both responders | Naïve | 1/1 | 1/1 | 1/1 |
|  | I/R Control | 9/10 | 9/10 | 8/10 |
|  | I/R | 3/3 | 2/3 | 2/3 |
|  | PenCON+I/R | 6/7 | 7/7 | 6/7 |
| | Penα1+I/R | 0/0 | 0/0 | 0/0 |

### P2X5 plays an important role in the development of pain related behaviors and increased EPRs after I/R

qPCR results suggested that GFRα1 regulated the development of muscle pain-related behaviors and exacerbated EPRs after I/R through modulation of the expression of ASIC3 and/or P2X5. Since we previously have shown that ASIC3 is crucial for I/R-related peripheral sensitization^16^, here we assessed the role of P2X5 on these phenomena. Using similar strategies to that described for GFRα1 knockdown, we confirmed that injection of Penetratin-linked P2X5 targeting siRNAs into the median and ulnar nerves of mice with I/R was able to completely block the I/R induced upregulation of this channel at the mRNA (I/R: 70%±10%; p<0.01 vs naïve; PenCON+I/R: 70%±20%; p<0.01 vs naïve; PenX5: −65%±36%;p>0.001 vs naïve; 1-way ANOVA with HSD post hoc) and protein levels (Fig 4A). Behavioral testing revealed that animals injected with P2X5-targeting siRNAs (PenX5+I/R) had lower guarding scores than both I/R and PenCON+I/R treated animals, although this did not reach sham levels (Fig. 4B). Reduced mechanical withdrawal thresholds to von Frey filament stimulation (Fig. 4C) were partially reversed by P2X5 knockdown after I/R. However, the reduction in muscle withdrawal thresholds to forepaw muscle squeezing (Fig 4D) were not significantly inhibited in the PenX5+I/R group. Grip strength deficits were partially rescued from the effect of I/R by PenX5 injection (Fig. 4E). Interestingly, PenX5 injection completely prevented the exacerbation of the EPR after I/R, similar to that observed with Penα1 injection. Results suggest that P2X5 (in possible conjunction with other receptors like ASIC3 ^16^), plays an important role in the development of pain related behaviors and enhanced EPRs after I/R.

**Figure 4.**
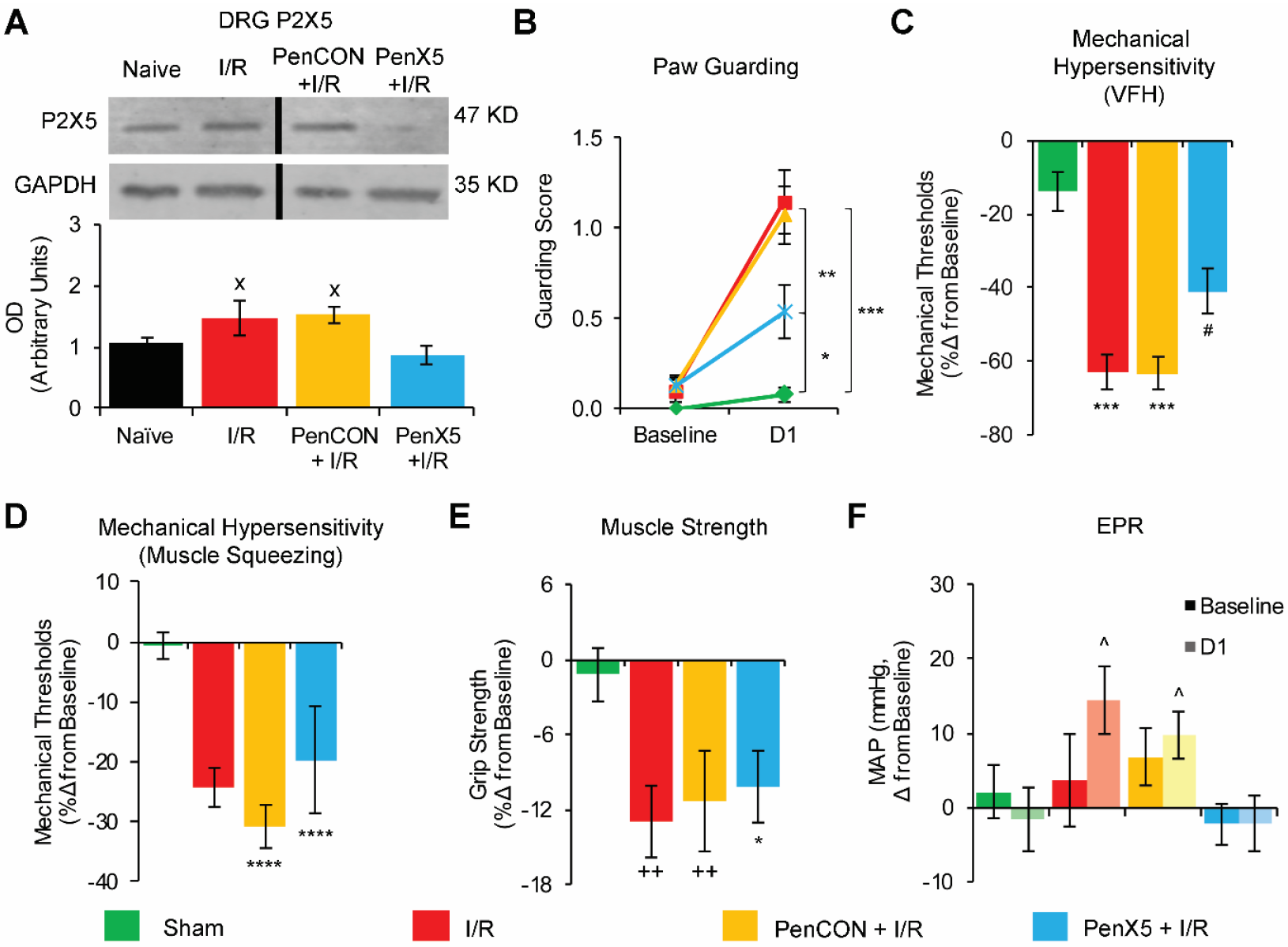
Selective knockdown of P2X5 in primary muscle afferents prevents the development of pain-related behaviors and exacerbated EPRs after I/R. **A**. P2X5 is increased in DRGs 1d after I/R or PenCON+I/R and this is prevented by PenX5 injection in mice with I/R (n=3 per group). **B.** Paw guarding is increased 1d after I/R and PenCON+I/R compared to sham controls, and this is partially prevented in PenX5+I/R animals. **C.** Mechanical withdrawal thresholds to von Frey hair (VFH) stimulation are decreased after I/R and PenCON+I/R and this is partially recued in the PenX5+I/R group. **D.** Mechanical withdrawal thresholds to muscle squeezing are decreased after I/R, PenCON+I/R and PenX5+I/R. **E.** PenX5 injection in mice with I/R also partially inhibited I/R-related grip strength deficits observed in I/R alone or PenCON+I/R groups. **F.** A significant increase in mean arterial pressure (MAP) after exercise was observed in injured mice (I/R and PenCON+I/R), but this was not observed in the PenX5+I/R group. (D-E n=12 per group. ^x^p<0.5 vs Naïve, *p<0.5 vs Sham, **p<0.01 vs PenX5+I/R, ***p<0.001 vs Sham and PenX5+I/R, ****p<0.0001 vs Sham, ++p<0.01 vs Sham, ^^^p<0.05 vs pre-exercise MAP; 1-way ANOVA with HSD post hoc test (A) or 2-way RM ANOVA with HSD post hoc (B-F)).

### CREB binding protein (CBP) inhibition prevents I/R induced overexpression of P2X5 and the development of ischemic-myalgia-like behaviors

Results suggested that enhanced GDNF/GFRα1 signaling increased P2X5 (and ASIC3) expression in muscle afferents to dually modulate nociception and EPRs after I/R. However, we did not know how GDNF signaling influenced transcription. We therefore performed RT array analysis of DRGs from mice with I/R to perform an unbiased screen of several transcription factors simultaneously. Surprisingly, no transcription factors were upregulated at the mRNA level in the DRGs 1d after I/R using this approach (Supplementary Table 1). We therefore retrogradely labeled afferents from the forepaw muscles using fluorogold, dissociated the DRGs *in vitro* and treated them with GDNF. Unlike our previous reports analyzing IL1β treated muscle afferents ^16^, GDNF was not found to alter the numbers cells containing activated c-Jun-N-terminal kinase (p-JNK). GDNF also did not increase the numbers of cells with phosphorylated mitogen activated protein kinase (MAPK) p38 (Supplementary Fig 1).

Since our previous reports suggested that MAPKs such as extracellular signal related protein kinases 1/2 (ERK1/2) were not upregulated or activated by I/R in the DRGs (21), we performed additional WB analysis from I/R injured animals to assess other putative transcription factors that may be activated (phosphorylated) at the protein level by increased GDNF/GFRα1 signaling. Although no changes in pERK5 were detected (not shown), we found that cAMP response element binding (CREB) protein was phosphorylated after I/R in the DRGs. However, Penα1 injection did not fully prevent I/R-induced CREB phosphorylation (Fig. 5A). Since CREB has been linked to receptor tyrosine kinase signaling ^45, 46^, and these molecules execute their functions through transcription factor complexes ^47, 48, 49^, we decided to assess the CREB binding protein (CBP) in mice with I/R. Not only did I/R upregulate CBP along with activation of CREB, but knockdown of GFRα1 in the DRGs after I/R prevented CBP upregulation (Fig. 5A).

**Figure 5.**
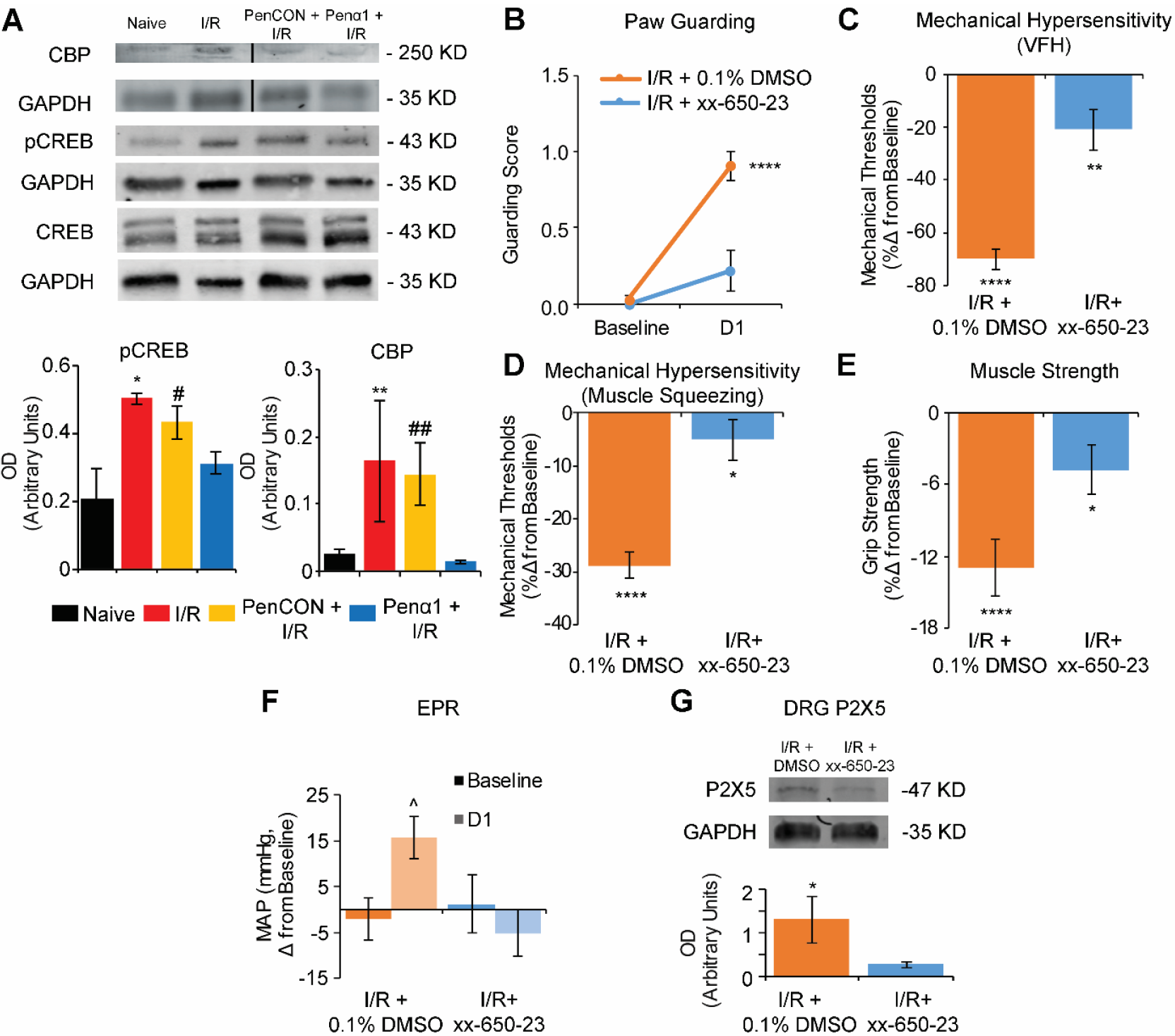
Disruption of the CBP/CREB transcription factor complex regulates I/R-related pain-like behaviors, EPRs and P2X5 expression. **A**. pCREB and CBP are increased after I/R compared to naïve as assessed with western blotting. Penα1 injection however partially blocked the I/R-induced upregulation of pCREB and fully inhibited the upregulation of CBP in the DRGs. **B.** Paw guarding is increased 1d after I/R in vehicle treated (0.1% DMSO) animals and this is completely prevented by treatment with the CREB-CBP interaction antagonist xx-650-23. Mechanical withdrawal thresholds to VFH stimulation **(C)** and muscle squeezing **(D)** are significantly decreased after I/R+0.1%DMSO and I/R+xx-650-23. However, both of these I/R-induced behavioral effects were significantly blunted in xx-650-23 treated animals. **E**. Decreased grip strength observed 1d after I/R+0.1%DMSO was partially rescued by xx-650-23. **F.** Exercise pressor reflexes (reflected by changes in MAP after forced running) exacerbated by I/R were also blocked by disruption of the CBP/CREB complex **G.** xx-650-23 significantly reduced the levels of P2X5 in the DRGs of mice with I/R compared to I/R-injured animals treated with vehicle (0.1% DMSO). (A: n=4-7 per group, *p<0.5 vs naïve but p<0.13 vs Penα1, #p<0.08 vs naïve but p<0.4 vs I/R, **p<0.05 vs naïve and Penα1, ##p<0.14 vs naïve and Penα1 but p<0.9 vs I/R; B-F: n=10 per group. *p<0.5 vs Baseline, **p<0.01 vs Baseline, ****p<0.0001 vs Baseline, ^^^p<0.05 vs pre-exercise MAP; G: n=3 per group, *p<0.05 vs I/R+xx-650-23; 1-way ANOVA with HSD post hoc test (A, G) or 2-way RM ANOVA with HSD post hoc test (B-F)).

We therefore used a pharmacological approach to assess whether disruption of the CBP/CREB transcription factor complex could blunt I/R-related hypersensitivity and altered EPRs. Treatment of mice with the CBP antagonist, xx-650-23 not only inhibited the ischemic-myalgia-like behaviors after I/R (Fig. 5B-F), but it also reduced the levels of P2X5 in the DRGs (Fig. 5G). Results indicated that I/R related-behaviors modulated by increases in GDNF/GFRα1 signaling, regulates P2X5 induction at least in part through modulation of CREB/CBP-mediated transcription.

### Targeting GDNF directly in the injured muscles effectively prevents the development of pain-related behaviors and exacerbated EPRs after I/R

To test whether targeting increased levels of muscle GDNF directly could prevent ischemic myalgia-like behaviors, we then injected either anti-GDNF antibodies, vehicle or IgG into the affected forepaw immediately after sham or I/R and assessed pain-related behaviors and EPRs one day after injury. Mice injected with vehicle or IgG were not found to be different from each other (not shown). However, as indicated for GFRα1, P2X5 and CBP inhibition, GDNF antibody injection into the muscles significantly inhibited I/R-induced paw guarding, mechanical hypersensitivity, grip strength and altered EPRs compared to control antibody injected mice with I/R (Fig. 6).

**Figure 6.**
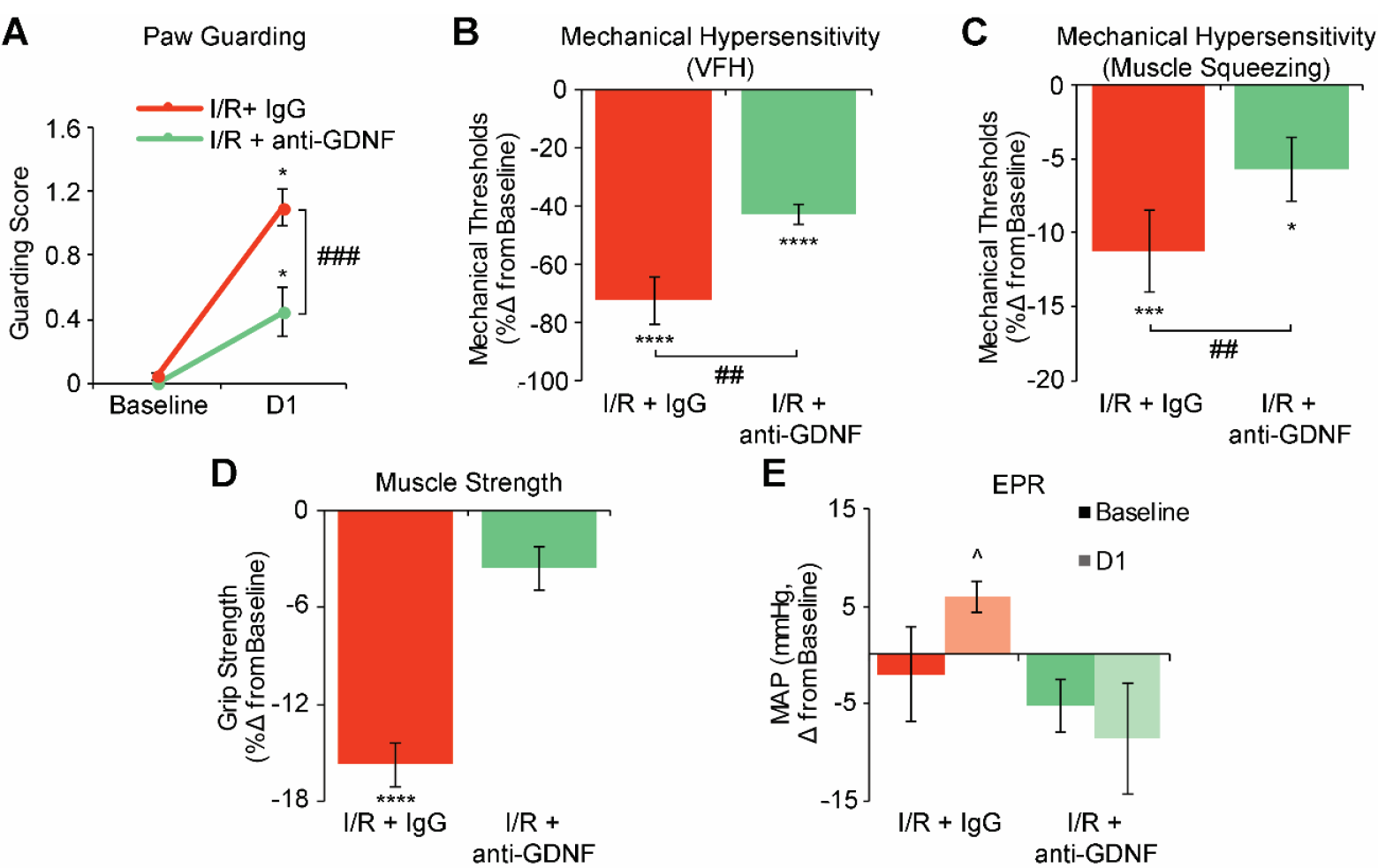
Intramuscular injections of anti-GDNF antibody partially prevents the development of pain related behaviors and enhanced EPRs after ischemia/reperfusion (I/R). **A**. Paw guarding is increased 1d after I/R+IgG and this is partially prevented by treatment with anti-GDNF at the time of injury. **B**. Mechanical withdrawal thresholds to VFH stimulation are significantly decreased after both I/R+IgG and I/R+anti-GDNF. **C**. Mechanical withdrawal thresholds to muscle squeezing are also decreased after I/R+IgG and I/R+anti-GDNF; however, anti-GDNF partially reversed I/R-related effects on muscle hypersensitivity. **D.** Muscle strength was decreased 1d after I/R+IgG but anti-GDNF injection completely prevented the reduction in grip strength. **E.** Changes in MAP after forced running were unaffected by IgG injection in mice with I/R but anti-GDNF treated animals did not show any changes in post exercise MAP. (A-E n=10 per group. *p<0.5 vs Baseline, **p<0.01 vs Baseline, ****p<0.0001 vs Baseline, ^##^p<0.01 vs I/R+anti-GDNF, ^###^p<0.01 vs I/R+anti-GDNF, ^^^p<0.05 vs pre-exercise MAP; 2-way RM ANOVA with HSD post hoc test).

## Discussion

In this report, we describe a novel mechanism of muscle afferent sensitization after ischemia with reperfusion injuries that regulate the development of pain-related behaviors and exacerbated cardiovascular responses to exercise. Peripheral sensitization after I/R was modulated by increased GDNF from injured muscles acting upon upregulated GFRα1 in DRG neurons which (among other possible players) modulated CREB/CBP dependent transcription of P2X5 (Fig. 7).

**Figure 7.**
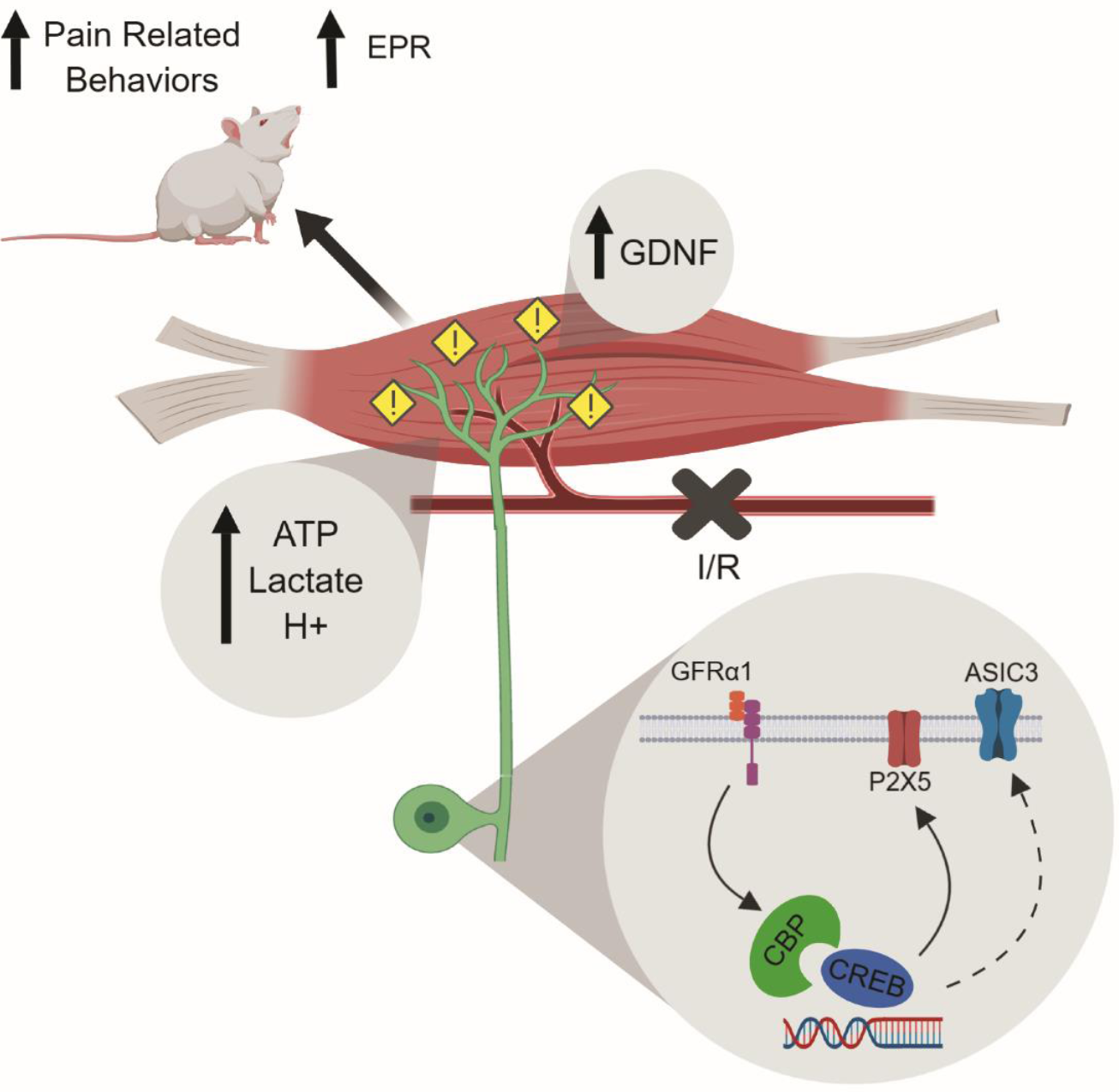
A role for peripheral GDNF signaling in the development of ischemic myalgia. After I/R, injured muscles release a variety of metabolites that include ATP, lactate and protons that stimulate group III and IV muscle afferents. Simultaneously, GDNF is being release in the injured tissues and this induces a signaling cascade in primary sensory neurons through its receptor GFRα1 that involves increased function of CREB/ CREB binding protein (CBP), which in turn modulates the observed overexpression of ATP sensitive channel P2X5 and to a lesser degree, ASIC3. This molecular change at the DRG level in turn modulates the appearance of pain-related behaviors such as paw guarding, mechanical hypersensitivity and muscle weakness. In addition, this pathway also affects the exercise pressor reflex after injury. Targeting the overexpression of P2X5 or GFRα1 in sensory neurons or the increased GDNF levels within the muscle appears to inhibit many of these ischemic myalgia-like phenomena.

Multiple models of musculoskeletal pain have suggested a prominent role for GDNF signaling ^32, 33, 34^. Various studies have shown that chemosensitive group III and IV muscle afferents are the sensory arm of the exercise pressor reflex and that ischemic injuries are capable of increasing their response to metabolite stimulation and muscle contraction ^5, 8, 14, 50^. In addition, group III and IV primary muscle afferents also function as nociceptors and can be sensitized by ischemic injury. GDNF/GFRα1 signaling is likely to be a major aspect of I/R-related hypersensitivity. Of all combinations of growth factors and their receptors upregulated in the muscles and DRGs, only GDNF and GFRα1 were collectively induced after I/R ^15^ (Table 1). Selective knockdown of GFRα1 in muscle afferents partially prevented injury-related paw guarding and completely blocked the I/R-induced decrease in mechanical withdrawal thresholds to VFH stimulation and to muscle squeezing (Fig. 1).

Interestingly, GFRα1 knockdown was only partially effective in preventing the decreased grip strength observed after injury suggesting that there are other mechanisms involved in altering specific muscle-related tasks after I/R such as cytokine-mediated regulation of ASICs ^16^. Previous work from our lab showed that preventing the I/R-induced upregulation of IL1r1 in DRGs completely prevented ASIC3 induction and in turn fully prevented the development of all pain-related behaviors after injury ^16, 17^. Studies using ASIC3 knockout mice report a lack of mechanical hyperalgesia after muscle inflammation that can be restored by expressing ASIC3 in the neurons innervating the affected tissue ^21, 22^. However, in this report, selective knockdown of GFRα1 fully blocked the I/R induced upregulation of the purinergic receptor P2X5, but only partially that of ASIC3. Together this provides a plausible explanation as to why Penα1 injection only partially blunted I/R-induced paw guarding and injury-reduced grip strength. Thus, pain-related behaviors and altered EPRs from I/R are due to activation of several signaling pathways that likely includes both cytokine and growth factor signaling mechanisms.

At the sensory neuron level, GFRα1signaling appeared to play a role in the development of primary afferent mechanical hypersensitivity after I/R, as GFRα1 knockdown blocked I/R induced reductions in group III/IV muscle afferent thresholds (Fig. 2). In addition, we observed a phenotypic switch from observing mostly two mutually exclusive sub-populations of chemosensitive fibers, metaboreceptors (low metabolite responders) or metabo-nociceptors (high metabolite responders), to a subpopulation that became responsive to both “low” and “high” metabolite concentrations. This finding, confirms our previous reports of a phenotypic switch after I/R ^15, 16^, but in contrast with this earlier study, selective knockdown of GFRα1 after I/R did not revert the phenotypic switch. Instead, it vastly decreased the numbers of detectable chemosensitive neurons, suggesting that GFRα1 signaling, especially after I/R, plays an important role in the maintenance of peripheral chemosensitivity.

This result could be explained by the downstream effects on P2X5 (Figs. 3-5), a key player in the chemosensitive function of muscle afferents ^7, 19^. P2X5 has been shown to modulate the pH sensitivity of ASIC3 *in vitro* ^19^. Previous work showed exposure to a combination of metabolite stimuli is necessary to effectively activate chemosensitive afferents ^1, 7^. Thus, it is reasonable to suggest that disrupting one of the components of the chemosensitive apparatus could render the primary neurons incapable of responding to a chemical stimulus (depending on the concentrations or combinations of metabolites present) ^7^. This could also explain why Penα1+I/R mice and PenX5+I/R mice do not exhibit an increased EPR after injury as chemosensitivity of group III/IV muscle afferents is a key component of EPR function ^6, 8, 11, 24, 51, 52^. Nevertheless, regulation of P2X5 expression via CREB/CBP dependent transcription (Fig. 6) may be a key means through which GFRα1 modulated muscle afferent sensitization after I/R.

An important consideration is that not all chemosensitive neurons are the same regarding GFRα1 or P2X5. Our neurochemical analysis of functionally identified chemosensitive neurons shows that metaboreceptors (“low” metabolite responders) may only partially rely on GFRα1 or P2X5 as only ∼33-25% (Table 2) of these cells co-expressed both receptors. In contrast, metabo-nociceptors (high metabolite responders), often express both GFRα1 and P2X5 (Fig. 4). After I/R, the new subpopulation of afferents that responds to both metabolite concentrations almost always co-expresses GFRα1 and P2X5, reinforcing the idea that enhanced co-expression of both receptors is important in the observed phenotypic switch in chemosensitive neurons after I/R.

The behavioral experiments on P2X5 targeted knockdowns somewhat support this notion, as the guarding scores on PenX5+I/R mice almost perfectly mimic those observed in the Penα1+I/R group. Yet, both mechanical hypersensitivity and muscle function were not rescued by PenX5 injection, suggesting that this aspect of afferent sensitization may be explained by concurrent cytokine/ ASIC signaling ^16^. Nevertheless, P2X5 knockdown in I/R affected afferents was quite effective in preventing exacerbated EPRs one day after injury, highlighting the importance P2X5 in modulating the chemosensitive function of muscle afferents. This complements previous *in vitro* reports that showed P2X5 can modulate ASIC3 pH sensitivity and could explain the increased behavioral and cardiovascular responses observed after injury here ^19^. Future studies are thus needed to assess afferent sensitization in mice with I/R and P2X5 knockdown.

Despite these novel findings, the origin of the observed increase in muscle GDNF levels after I/R is still up for debate. Our results suggested that GDNF could come directly from the I/R-affected myofibers (Fig. 1). Previous studies support this notion in that skeletal myofibers themselves are capable of releasing GDNF ^53, 54^. The immunohistochemistry performed in injured muscles also shows a distinct “halo-like” pattern around the injured muscle cells that highly resembles previous reports hinting that GDNF could originate in myocytes ^54^. While the effect of GFRα1 upregulation in the DRG after I/R appears to be an important player in injury-related hypersensitivity, the cause of increased receptor expression was not elucidated. Additional studies are also needed to determine how GFRα1 is initially upregulated after injury.

The observed increase in MAP after I/R resembles previous reports in other models of ischemic insult ^13, 14, 52, 55, 56^. However, this is the first report where *unilaterally* targeting a neurotrophic factor or its receptor, prevented the development of exacerbated EPRs post injury. While we were able to detect an exacerbated MAP after I/R, we did not observe significant changes in heart rate (HR) after exercise before or after injury. This could be due to a fast recuperation of HR after exercise compared to the observed MAP changes. Nonetheless, the fact that our strategies targeting GDNF/GFRα1 were effective at preventing the exacerbated EPR after I/R reveal how important this signaling pathway may be for the sensitization of muscle afferents and subsequent effects on cardiovascular reflexes or nociceptive-like responses.

### Clinical Significance

Patients that experience pathologies characterized by musculoskeletal ischemic injury, such as sickle cell disease, PVD, or complex regional pain syndrome (CRPS) often present both pain and altered autonomic vasomotor responses to tissue ischemia ^36, 57, 58, 59^. Our data suggests that muscle GDNF, combined with GFRα1 upregulation can sensitize muscle primary afferents after I/R. To determine whether targeting this pathway was a potential therapeutic for I/R-related hypersensitivity, we targeted GDNF at the source by injecting antibodies into the I/R injured muscles similar to strategies used in other disease models ^60, 61, 62^. Our results highlight the potential therapeutic value of blocking this pathway at the source in ischemic injuries as our mice displayed reduced pain-related behaviors and EPRs after I/R upon intramuscular injection of GDNF targeting antibodies (Fig. 7). Results thus suggest that reducing GDNF signaling may be an option to treat patients that suffer musculoskeletal ischemic injuries and that this strategy may not only alleviate their pain, but also prevent cardiovascular complications associated with this injury.

## Methods

### Animals

Experiments were conducted with young adult (3–8 wks) male Swiss Webster mice. For all cohorts of the behavioral experiments involving anti-GDNF injections or xx-650-23 injections (and controls), mice were purchased from Envigo (Madison, WI). For experiments involving siRNA injections, animals were obtained from Charles River (Wilmington, MA). No baseline differences were found between mice obtained from these two vendors (not shown). All animals were housed in a barrier facility in group cages of no more than 4 mice, maintained on a 12:12-h light-dark cycle with a temperature-controlled environment, and given food and water ad-libitum. All procedures were approved by the Cincinnati Children’s Hospital Research Foundation Institutional Animal Care and Use Committee and adhered to NIH Standards of Animal Care and Use under AAALAC approved practices.

### Induction of ischemia with reperfusion injury

Ischemia with reperfusion injury (I/R) was induced as previously described ^15^. Briefly, mice were anesthetized with 3% isoflurane, and then under a dissection microscope, a small incision was made on the right forelimb above the elbow. The right brachial artery was exposed proximal to the ulnar artery/radial artery split. The vessels were gently loosened from adjacent connective tissue and then the brachial artery was tied using a 7-0 silk suture using a loop knot. Incisions were closed with 6-0 silk sutures, and six hours after performing the occlusion surgery, a second surgery was performed to remove the suture around the artery. Except for the baseline behavioral time points, all assessments were performed 18 hours after the suture removal (reperfusion). For comparisons, some mice received a sham I/R surgery in which a suture was placed under the artery as described above, but it was not tied.

### Nerve-targeted siRNA injections

Specific targeting siRNAs were used to selectively knockdown the expression of either the GDNF receptor, GFRα1, or the ATP receptor, P2X5 (Thermo). siRNAs were conjugated to Penetratin-1 (MP Biomedicals) as previously described ^13, 43, 44, 63^. The duplex used for each target in these experiments was determined to have the highest targeting efficiency based on the knockdown efficacy of four different targeting siRNAs tested in Neuro2A cells *in vitro* (not shown). The duplex used in the current study that was found to most efficiently target GFRα1 was as follows: sense: 5’-S-S-CGACAAAGUUCCAGCCAAGUU; anti-sense: 5’-P-CUUGGCUGGAACUUUGUCGUU. The duplex used to target P2X5 was: sense: 5’-S-S-CAACAUUGGUUCCGGGCUGUU; anti-sense: 5’-P-CAGCCCGGAACCAAUGUUGUU. The non-targeting control siRNA used this study is the same as in previous work from our lab ^13, 16, 44^, and has been determined to not target any gene in the mouse genome (Thermo; Cat#: D-001206-14-05).

Two days before I/R, mice were anesthetized as described. A small incision was made in the inner mid-forelimb region, proximal to the elbow and exposed the ulnar and median nerves to be injected. siRNAs were heated to 65°C for 5 min prior to injection. 0.1–0.2 μL of 90 μM penetratin-1 linked non-targeting, control (PenCON), GFRα1-targeting (Penα1), or P2X5-targeting siRNAs (PenX5) were pressure injected directly into the median and ulnar nerves using a quartz microelectrode connected to a picospritzer. This procedure does not cause significant injury to the sensory neurons being studied and does not it induce antiviral related responses ^16, 41, 43, 44^.

### Antibody injections

Immediately after the occlusion surgery and while still under isoflurane anesthesia, cohorts of mice were slowly injected with 10µg of either anti-GDNF, antibody (ANT-014, Alomone labs, Jerusalem, Israel), IgG (AB-105-C, R&D Systems, Minneapolis, MN, USA) or vehicle (sterile water) in a volume of 5µL into the right forepaw muscles using a 30 gauge needle. Antibody dose was determined from Murase et al (45). Reperfusion surgery in these mice was performed 6h later as described above.

### CREB binding protein (CBP) antagonist injections

The compound xx-650-23 was used to block the interaction between the cAMP response element-binding (CREB) protein and CREB binding protein (CBP). 20 mg/kg of xx-650-23 (GLXC-12046, Glixx lab, Hopkinton, MA, USA) or an equivalent volume of vehicle (0.1% DMSO) was injected intraperitoneally (ip.) in a volume <200uL immediately after occlusion surgery under isoflurane anesthesia. Pilot dose-response experiments in which small cohorts of animals were treated with either 10, 20 or 40mg/kg xx-650-23 were used to determine the dose used in these experiments (20mg/kg). Mice were allowed to recuperate after the injection and reperfusion followed 6h later.

### Measurement of pain-related behaviors and exercise pressor reflexes (EPRs)

Separate groups of sham, I/R, siRNA+I/R (PenCON+I/R, Penα1+I/R or PenX5+I/R), local antibody injection+I/R (anti-GDNF+I/R, IgG+I/R or vehicle+I/R), or antagonist+I/R (xx-650-23+I/R or 0.1% DMSO+I/R) were used for behavioral analysis (n = 10 per group). Testing of pain-related behaviors was performed as previously described by Ross et al., 2014. Briefly, mice were first tested at baseline, one day prior to injury (I/R), and then again one day post injury. All behavioral testing was performed in the morning. The experimenter was blinded to both treatment and injury condition of the animal.

Nociceptive testing included 4 behavioral assessments: forepaw guarding, von Frey filament stimulation of the plantar surface of the forepaws, and forelimb grip strength, in this order. Fore paw muscle squeezing was performed in independent cohorts (below). Mice were placed in a raised acrylic glass chamber with a steel mesh bottom and allowed to habituate for at least 30 minutes. To evaluate guarding behavior, mice were assigned a score of 0 to 2 (0 = mouse places foot firmly on mesh, 1 = mouse does not bear full weight on foot, 2 = mouse holds foot completely above mesh) every 5 minutes for 12 total observations. The average score for the 12 trials was determined for each mouse per behavioral day. Mechanical withdrawal thresholds on the forepaws were then determined by stimulating the plantar surface with an increasing series of von Frey filaments (0.07 g to 4 g). Threshold to withdrawal was recorded for at least 3 trials with 5-minute intervals between stimulations, and the average of the 3 trials was used for analysis. Mice were then assayed for forepaw muscle strength using a grip strength meter (BioSeb, Vitrolles, France). Animals were held by the tail over a metal grid until they firmly held it with both fore paws but were not allowed to grip the grid with their hind paws. Then they were quickly pulled back horizontally (along the axis of the force sensor) until they could not retain their grip. Grip strength was measured (in g) in 3 rounds of 3 trials each, with 5 minutes between each round. The average of the nine trials was used for analysis. Finally, cohorts of mice were tested using a digital Randall-Sellito device (IITC Life Science Inc. Woodland Hills, CA, USA) to assess withdrawal thresholds to muscle squeezing. In order to diminish the stimulus applied to the skin and maximize the stimulus to the muscle we used a blunt, rounded probe ∼1.5mm wide, similar to approaches previously described ^64, 65^. To prevent injury due to excessive pressure, a cutoff pressure was set to 400g. Animals were subjected to 3 trials with an interval of 5 minutes between each test. Average of the three trials was used for analysis.

After all pain-related behavioral assessments, the exercise pressor reflex (EPR) was then determined by using a low intensity, forced run protocol based on previous work by our lab and others ^13, 66, 67^. Before and immediately after the exercise session, each mouse was placed in a small acrylic restrainer adequate for its size and weight, allowed a short period of acclimation (approximately 5 minutes) and had its blood pressure and heart rate measured 45 times (maximum number obtained per mouse) with a tail cuff BP system (Kent Scientific, Torrington, Connecticut, USA). Data was collected using CODA software (Kent Scientific) and analyzed offline. The first 5 measurements were used to acclimate the mice to tail cuff inflation and were not used for analysis. Unreliable measurements were automatically discarded by the software and manually when the animal showed excessive movement that generated signal artifacts.

For the exercise protocol, the mice were run on a modular treadmill (Columbus Instruments, Columbus Ohio) at 0° of inclination with an increasing ramp speed going from 9 m/min up to a maximal speed of 13 m/min for a total distance of 500m. Speed was increased 1 m/min per minute, thus the entire running protocol lasted approximately 40 minutes. This speed is around 75% the mean critical speed for mice and is well below the speed and distance previously reported to induce anaerobic metabolism in the muscle or tissue damage in mice ^13, 68, 69^. This exercise protocol has also not been shown to induce pain-related hypersensitivity due to ischemic injury ^13^.

### *Ex vivo* recording preparation

*Ex vivo* recording was performed as previously described by Jankowski et al (2013) and Ross et al (2014). Briefly, mice were anesthetized with an intramuscular hindlimb injection of ketamine and xylazine (100 and 16 mg/kg, respectively) and perfused transcardially with oxygenated (95% O2-5% CO2) artificial cerebrospinal fluid (aCSF; in mM: 127.0 NaCl, 1.9 KCl, 1.2 KH2PO4, 1.3 MgSO4, 2.4 CaCl2, 26.0 NaHCO3, and 10.0 D-glucose) at 12–15°C. The right forelimb and the spinal cord (SC) were then excised and placed in a bath of the same aCSF. The forelimb skin was removed along with the cutaneous branches of the median and ulnar nerves. SC was hemisected and the median and ulnar nerves along with the forelimb muscles they innervate (with bone left intact) were dissected in continuity with their respective DRGs (C7, C8, and T1). After dissection, the preparation was transferred to a separate recording chamber containing cold, oxygenated aCSF. The forepaw was pinned on an elevated platform, keeping the entire paw perfused in a chamber isolated from the DRGs and the SC. Finally, the bath was slowly warmed to 32°C before recording from the DRGs.

All single unit recordings were made from the C7, C8, and T1 DRGs as these are the primary source of muscle afferent fibers in the median and ulnar nerves. Sensory neuron somata were impaled with quartz microelectrodes (impedance>150MΩ) containing 5% Neurobiotin (Vector Laboratories, Burlingame, CA) in 1M potassium acetate. Electrical search stimuli were delivered through a suction electrode on the nerve to locate sensory neuron somata with axons in the median and ulnar nerves. The latency from the onset of this stimulus and the conduction distance between the DRG and the stimulation site (measured directly along the nerve), were used to calculate the conduction velocity (CV) of the fibers. Group IV afferents were classified as those with a CV≤1.2 m/s, and group III afferents were those with CVs between 1.2 and 14 m/s ^1, 13^. Peripheral receptive fields (RFs) in the muscles were localized by electrically stimulating the muscles with a concentric bipolar electrode. Only driven cells with RFs in the muscles then underwent mechanical, thermal, and chemical testing. Mechanical response characteristics were assessed with an increasing series of von Frey hairs ranging from 0.4 g to 10 g (with diameters of 0.23–0.36 mm). Mechanical stimulation of the RF was held for approximately 1-2 seconds. Thermal responses were determined by applying hot (≥52°C) or cold (≤3°C) saline directly to the paw muscles at the electrically determined RF. Each application lasted ∼1-2s. After that, the muscles were exposed to an oxygenated “low” metabolite mixture (15 mM lactate, 1 µM ATP, pH 7.0) and then to a “high” metabolite mixture (50 mM lactate, 5 µM ATP, pH 6.6) delivered by a valve controller with an in-line heater to maintain solutions at bath temperature. ATP was added to the mixture immediately prior to delivery of metabolites. Adequate recovery times (∼20–30 s) were employed between all stimulations. Elicited responses were recorded digitally for off-line analysis (Spike2 software, Cambridge Electronic Design). After physiological characterization, select cells were labeled by iontophoretic injection of 5% Neurobiotin (1 or 2 cells/DRG).

### Immunohistochemistry

Once a sensory neuron was characterized and intracellularly filled with Neurobiotin, the DRG containing the injected cell was removed and immersion fixed with 3% paraformaldehyde in 0.1 M phosphate buffer (PB). DRGs were fixed for 30 min at room temperature (RT) and then changed to PB for storage before embedding in OCT embedding medium. Embedded DRGs were stored at −80°C, sections (15 µm) were cut on a cryostat and mounted on slides and processed for GDNF receptor GFRα1 (goat anti-GFRα1, 1:100; cat no. AF-560, R&D Systems, Minneapolis, MN, USA) and P2X5 (rabbit anti-P2X5, 1:100; cat no. APR005 Alomone Labs, Jerusalem, Israel). After incubation in primary antiserum, sections were washed then labeled with secondary antibodies (anti-goat AlexaFluor 568 cat. No. A31573, 1:400; or anti-rabbit AlexaFluor 647, 1:400; Thermo Fisher Scientific, Waltham, MA, USA) as well as FITC-Avidin (cat. No. A-2001, 1:700; Vector Laboratories, Burlingame, CA, USA) to label Neurobiotin-filled cells. Sections were mounted with Fluoromount G (cat. No. 17984-25, Electron Microscopy Sciences, Hatfield, PA) and stored in the dark at room temperature. Exposure time during microscopic analysis for each positive and negative image was performed at the same intensity level to confirm staining above background. Distribution of fluorescent staining was determined with a Nikon confocal microscope with sequential scanning to avoid bleed-through of the fluorophores. Images for publication were prepared using Photoshop Elements software (Adobe, San Jose, CA, USA).

### Quantification of DRG neurons

Total cells containing GFRα1 and P2X5 were quantified in DRGs (C7 or C8) from naïve mice, or mice that underwent I/R alone, PenCON+I/R, Penα1+I/R or PenX5+I/R (n=3 per condition). The DRGs were taken after electrophysiological experiments and were processed for immunohistochemical analysis as described above. The numbers of positive cells were determined using a slightly modified methodology as previously reported by Christianson et al (2006) and Jankowski et al (2009) to account for the thinner (15μm) sections obtained for immunocytochemical processing here. In brief, three non-consecutive groups containing three sections in series (45μm total) were randomly chosen and Z-stacks were generated at 3µm intervals to create 15µm thick optical sections using a Nikon confocal microscope with sequential scanning. The number of GFRα1-positive and P2X5-positive cells in each group were counted using Neurolucida software ensuring that the same cell was not counted twice including those in serial sections, averaged and reported as mean ± SEM.

### Western Blotting

C7-T1 DRGs were collected from the right side of sham injured, I/R, PenCON+I/R, Penα1+I/R, PenX5+I/R, xx-650-23+I/R or 0.1% DMSO+I/R mice (n=3 per condition). DRG tissue was pooled (2 mice of the same condition per sample). Forepaw muscle tissue was also collected from sham injured and I/R treated mice. All samples were collected one day after I/R in all groups. Tissue was homogenized in lysis buffer containing 1% sodium dodecyl sulfate (SDS), 10 mM Tris–HCl (pH 7.4), and protease inhibitors (1 µg/mL pepstatin, 1 µg/mL leupeptin, 1 µg/mL aprotinin, 1 mM sodium orthovanadate, and 100 µg /mL phenylmethylsulfonyl fluoride (Sigma-Aldrich Biochemicals, St. Louis, MO, USA). All samples (10 µg) were then centrifuged, boiled 10 minutes in 4X Protein Loading Buffer (cat. No. 928-40004, Licor), separated on either 12% (to quantify GAPDH, GFRα1, P2X5, ERK5, pERK5, CREB, pCREB protein), “any kD” (to quantify GDNF protein) or 7% (to quantify CBP) precast polyacrylamide gel (Bio-rad, Hercules, Cal, USA) and transferred to a polyvinylidene difluoride membrane (PVDF; Merck Millipore Ltd., Tullagreen, Ireland) that was blocked in Licor Blocking Buffer diluted 1:1 in 0.1M PB. Membranes were then incubated in this same blocking solution containing 0.2% tween-20 and primary antibodies overnight at 4°C. Antibodies used for western blot were as follows for each protein: GAPDH: chicken anti-GAPDH, 1:2000 (Pro-Sci, Atlanta, GA, USA); GFRα1: goat anti-GFRα1 1:250 (cat. No. AF560, R&D Systems, Minneapolis, MN, USA); P2X5: rabbit anti-P2X5 1:800 (cat. No. APR005, Alomone Labs, Jerusalem, Israel); GDNF: rabbit anti-GDNF 1:500 (cat No. ANT-014, Alomone Labs, Jerusalem, Israel); CREB: rabbit anti-CREB 1:1000 (cat. No. ab32515, Abcam, Cambridge, MA, USA); pCREB: rabbit anti-pCREB 1:2500 (cat. No. ab32096, Abcam, Cambridge, MA, USA); CBP: rabbit anti-CBP 1:500 (cat. No. ab2832, Abcam, Cambridge, MA, USA).

After incubation in primary antibodies, PVDF membranes were then washed and incubated with the appropriate secondary antibody (Donkey anti-chicken Dy680RD, 1:20000; Donkey anti-rabbit Dy800CW or Dy680RD, 1:20000; Donkey anti-goat Dy800CW, 1:200000. Licor, Lincoln, Nebraska, USA) diluted in blocking buffer containing 0.2% Tween-20 and 0.01%SDS. Membranes were washed and imaging was performed using a LICOR laser scanner for protein analysis. Immunoreactive bands were analyzed by measuring and plotting the band intensity and subsequently calculating the area under the curve using National Institutes of Health Image J software. Band intensity was normalized to GAPDH and reported as a ratio of gene/GAPDH expression. Phosphorylated proteins were then presented as a function of their total protein (e.g. pCREB/CREB or pERK5/ERK5).

### RNA isolation and reverse transcription, realtime PCR and transcription factor PCR arrays

DRG (C7-T1, right side) tissue was collected from cohorts of naïve, I/R, PenCON+I/R and Penα1+I/R mice. Separate cohorts of the same conditions were collected for transcription factor PCR arrays described below (n=3 per condition). RNA was isolated using the Qiagen RNeasy kit, according to the manufacturer’s protocol. For PCR arrays, samples were converted into cDNA following the RT^2^ First Strand Kit according to the manufacturer’s protocol. Then cDNA samples were mixed with an RT² Profiler™ SYBR Green PCR Array, realtime PCR master mix (Qiagen) and loaded into a RT² Profiler™ PCR Array Mouse Signal Transduction Pathway Finder™ (cat# PAMM-014Z, Qiagen). These plates were then run on an Applied Biosystems model StepOne Plus PCR machine.

For standard quantitative realtime PCR, 500ng of total RNA was DNAse I treated (Invitrogen) and reverse transcribed using Superscript II (Invitrogen) reverse transcriptase. 20ng of cDNA were used in SYBR Green realtime PCR reactions that were performed in duplicate and analyzed on a Step-One realtime PCR machine (Applied Biosystems). ASIC1, ASIC3, and GAPDH primer sequences (forward and reverse) were obtained from Ellit et al. ^70^ and GFRα2, GFRα3, trkA, trkB, and trkC obtained from Jankowski et al (51) for realtime PCR reactions. Primer sequences used for P2X3, P2X4, P2X5 and GFRα1 have also been detailed previously ^1, 15^.

Cycle time (Ct) values for all targets were all normalized to a GAPDH internal control. Ct values (used to determine fold change after injury) were then obtained by subtracting the normalized target gene’s Ct value from naive controls. Then fold change was determined as 2^Ct^ (Applied Biosystems). The error of the difference in means is then also calculated for the fold-change. Values were then converted and reported as a percent change where 2-fold change = 100% change.

### Retrograde labeling, DRG dissociation and GDNF treatments

To isolate afferents innervating the forepaw muscles, mice were anesthetized under isoflurane and 10μL of 4% fluorogold (FG; Fluorochrome) in 0.9%NaCl was injected into the right forepaw muscles using a syringe with 30g needle similar to that described previously (9). After 10d, mice were anesthetized and intracardially perfused with ice cold Hank’s balanced salt solution (HBSS). C7-T1 DRGs were isolated and collected in HBSS. DRG neurons were dissociated by first incubating the DRGs in cysteine/papain (0.03%, Sigma/ 20U/mL, Worthington) and then collagenase II (0.3%, Worthington). Digested DRGs were then triturated in F12 complete medium (media containing 10% fetal bovine serum and 1% penicillin/streptomycin) using fire polished glass Pasteur pipettes. Equal densities of cells were plated onto poly-D-lysine/laminin (20μg/mL each, Sigma) coated glass coverslips housed in 35mm petri dishes. Cells incubated at 37°C/5%CO_2_ for approximately 1hr prior to flooding the dishes with F12 complete media alone or complete F12 media containing 100ng/mL GDNF (R&D Systems). Cells incubated overnight at 37°C/5%CO_2_ prior to being rinsed in PB and fixed in 3% paraformaldehyde. Cells then underwent immunocytochemical processing similar to that described above using primary antibodies against activated c-Jun-N-terminal kinase (rabbit anti-pJNK, 1:100, Cell Signaling Technology) or phosphorylated p38 MAPK (rabbit anti-p-p38, 1:200, Cell Signaling Technologies) and appropriate secondary antibodies.

For quantification, five randomly selected fields from a given coverslip from each condition (n=5-7 per group) were imaged at 20x on a Leica inverted fluorescence microscope and analyzed offline using Adobe Photoshop. Cells were counted in each image and the total number of FG+, pJNK+ and/or p-p38+ cells were determined from the five fields per condition and averaged across conditions. Percentage of FG+ cells containing the marker of interest were reported.

### Statistics

All datasets were analyzed using Prism statistical software. Data was first assessed for normality and equal variance prior to performing the indicated statistical analyses. Specifically, WBs were analyzed by one-way analysis of variance (ANOVA, i.e. t-test) when two conditions were analyzed and one-way ANOVA with Holm-Sidak (HSD) post hoc test when multiple conditions were compared. For behavioral experiments, paw guarding was analyzed by two-way repeated measures (RM) ANOVA with Holm-Sidak post-hoc test for multiple comparisons. Mechanical hypersensitivity either to VFH or muscle squeezing, muscle strength and mechanical thresholds of primary afferents obtained with our *ex-vivo* preparation were analyzed using one-way ANOVA with Holm-Sidak post-hoc test or 1-way ANOVA/t-test when only 2 groups were being compared.

EPRs were analyzed using a two-way RM ANOVA with Holm-Sidak post-hoc test for multiple comparisons. The distribution of afferent subtypes in our groups obtained using *ex-vivo* recording was analyzed using a χ-square (χ2) test. DRG cell counts and RT qPCR data was analyzed either by one-way ANOVA with Holm-Sidak post-hoc test or 1-way ANOVA/t-test when only 2 groups were being compared. Analysis of cell counts *in vitro* from Supplementary Figure 1 were performed using Fisher’s exact test. Data are represented as mean ± SEM. Critical significance was defined with p≤0.05.

## Author contributions

Luis F. Queme performed most of the behavioral and molecular biology experiments, all of the electrophysiological recordings, collected tissue samples, analyzed the data and wrote the manuscript. Alex A. Weyler performed some behavioral experiments in addition to qPCR and Western Blot experiments. Renita C Hudgins performed some of the immunohistochemistry. Elysia R Cohen performed the cell culture experiments highlighted in supplementary figure 1. Michael P. Jankowski conceptualized and performed some of the experiments, structured and revised the manuscript and provided overall supervision of this work.

## Acknowledgments

This work was supported by grants to MPJ from the NIH/NIAMS (R01AR064551), and to LFQ from the American Heart Association (16POST29750004).

## Supplementary Materials

**Supplementary Figure 1.**
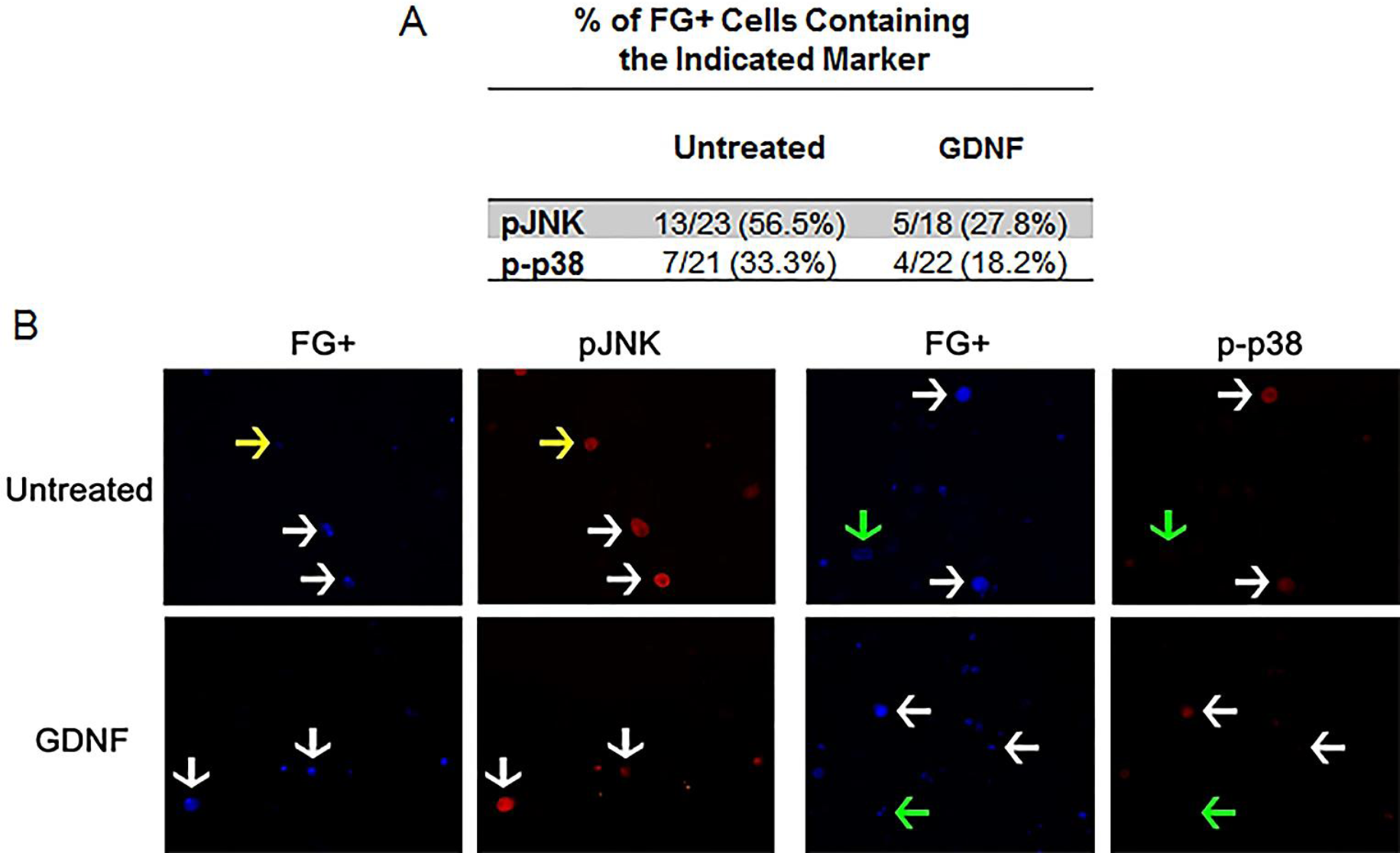
Treatment of retrogradely labeled muscle afferents with GDNF does not alter the numbers of cells expressing pJNK or p-p38. **A**: Quantification of fluorogold (FG) positive DRG cells *in vitro* treated with GDNF. p>0.05, Fisher’s exact test. **B**: Example images of FG+ cells (blue) immunostained for p-JNK or p-p38 (red). Yellow arrow: FG-/marker+; Green arrow: FG+/ marker-; White arrows: FG+/marker+.

**Supplementary Table 1:** RT array analysis of multiple transcription factors in the DRGs after I/R. Values indicate percent change versus naïve. mean ± SEM. No differences observed compared to naïve. p>0.05, 1-way ANOVA.

| Gene:<br>DRGs | I/R | Gene:<br>DRGs | I/R |
| --- | --- | --- | --- |
| Ar | 73 ± 31 | Mef2a | 66 ± 30 |
| Arnt | 60 ± 34 | Mef2b | -15 ± 28 |
| Atf1 | 61 ± 34 | Mef2c | 12 ± 33 |
| Atf2 | 68 ± 34 | Nfat5 | 70 ± 37 |
| Atf4 | 70 ± 27 | Nfatc2 | 49 ± 44 |
| Cebpa | 54 ± 35 | Nfatc3 | 65 ± 39 |
| Cebpb | 17 ± 36 | Nfatc4 | 67 ± 37 |
| Cebpg | 63 ± 32 | Nfkb1 | 51 ± 33 |
| Clasrp | 12 ± 34 | Nfyb | 75 ± 35 |
| Ctnnb1 | 46 ± 32 | Nr3c1 | 61 ± 33 |
| Dr1 | 59 ± 37 | Ppara | 49 ± 39 |
| E2f1 | 36 ± 27 | Pparag | 38 ± 44 |
| E2f6 | 58 ± 31 | Rb1 | 61 ± 32 |
| Egr1 | 56 ± 33 | Rel | 56 ± 36 |
| Esr1 | 51 ± 28 | Rela | 14 ± 40 |
| Ets1 | 45 ± 38 | Smad4 | 65 ± 35 |
| Ets2 | 52 ± 40 | Smad5 | 50 ± 37 |
| Fos | 52 ± 35 | Smad9 | 37 ± 32 |
| Gata2 | 62 ± 38 | Sp1 | 51 ± 39 |
| Gli1 | 31 ± 38 | Sp3 | 84 ± 33 |
| Gtf2b | 67 ± 33 | Stat1 | 67 ± 32 |
| Gtf2f1 | 56 ± 33 | Stat2 | 59 ± 32 |
| Htac1 | 44 ± 34 | Stat3 | 64 ± 35 |
| Hif1a | 54 ± 39 | Stat5b | 42 ± 33 |
| Hoxa5 | 47 ± 32 | Stat6 | 43 ± 35 |
| Hsf1 | 37 ± 30 | Tbp | 63 ± 32 |
| Id1 | 57 ± 30 | Tcf712 | 61 ± 34 |
| Irf1 | 36 ± 28 | Tfap2a | 39 ± 45 |
| Jun | 53 ± 33 | Tgif1 | 29 ± 41 |
| Jund | -7 ± 39 | Trp53 | 53 ± 34 |
| Kcnha | 59 ± 32 | Yy1 | 56 ± 34 |
| Max | 68 ± 30 |  |  |

## Notes

**Conflict of interest:** The authors declare no competing financial interests.

#### Summary of Updates

Updated title and abstract. Corrected formatting.

